# Isolation and functional characterization of the rat Thyroglobulin gene upstream enhancer region

**DOI:** 10.1101/146589

**Authors:** Christiane Christophe-Hobertus, Daniel Christophe

## Abstract

We report here the isolation and functional characterization of the as yet undescribed rat Thyroglobulin gene upstream enhancer element.

---

As shown in fig. 1, the alignment of the DNA sequences of the already characterized bovine (Christophe-Hobertus et al., 1992) and human (Berg et al., 1996) Thyroglobulin (Tg) gene upstream enhancer elements with genomic DNA sequences located upstream from the Tg gene transcribed region in mouse and rat identified the presence of four highly conserved sequence motifs. Among these were two stretches (boxed in fig. 1) that corresponded to the doublet of TTF-1 binding sites which had been shown to constitute the functional unit of the enhancer in the bovine (Christophe-Hobertus and Christophe, 1999a). It is noteworthy to point out here that the distance between the two TTF-1 binding sites is much shorter in both rodent species as compared to that in man and the bovine. In the bovine, a third TTF-1 binding had been identified in this place, but our previous work led to the conclusion that it is dispensable for enhancer function (Christophe-Hobertus and Christophe, 1999a). The absence of the corresponding DNA sequences in rodents thus further supports our previous conclusion at first sight.

**Fig. 1.**
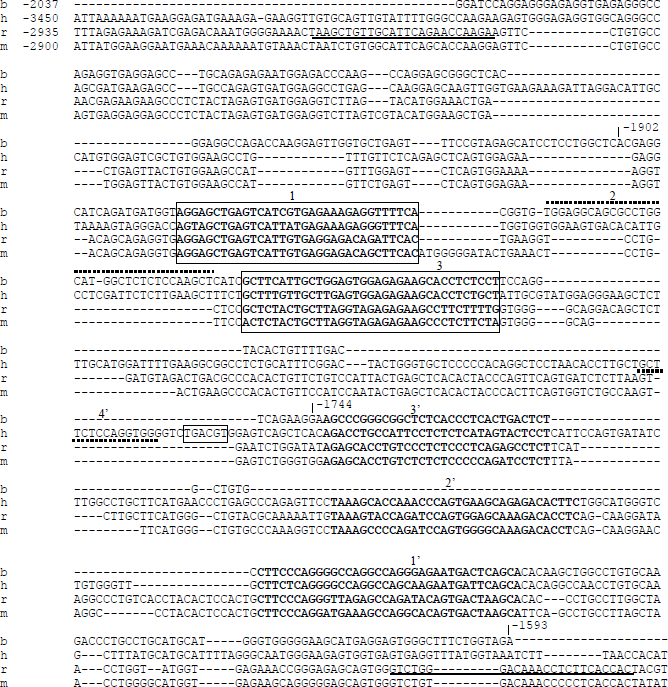
DNA sequences alignment of bovine (b) and human (h) enhancer elements with upstream regions from rat (r) and mouse (m) Tg gene. The position of the starting nucleotide relative to transcription start (+1) is given in the first line for each sequence. The boundaries of the DNaseI-hypersensitive enhancer region (−2037 to −1593) and core enhancer element (−1902 to −1744) identified previously in the bovine are also indicated. DNA sequences exhibiting the highest conservation scores (according to the DIALIGN 2.2.1 software used) are depicted in bold and numbered according to the corresponding TTF-1 footprints (see fig. 2). The sequences corresponding to the TTF-1 binding sites doublet that was shown to be critical for enhancer function in the bovine are boxed. The CRE-like element present in man only (boxed) and TTF-1 binding sites not found in the rodent sequences (dotted lines + numbers according to fig. 2) are also highlighted. Underlined sequences correspond to the primers used for PCR amplification of the rat enhancer region.

The two other conserved sequence motifs corresponded to TTF-1 footprints identified on the human enhancer sequence (Berg et al., 1996) that were contained in the DNaseI-hypersensitive enhancer region characterized initially in the bovine, but that were located outside of the core enhancer element as delimitated in the functional study of this enhancer region (Christophe-Hobertus et al., 1992). Another partially conserved motif was present in the human and rodent sequences only. It also corresponded to a known TTF-1 footprint on the human enhancer sequence (Berg et al., 1996). No specific function has been assigned to these DNA sequences as yet. A schematic representation of the available data on the organization of TTF-1 binding sites within rat, bovine and human Tg gene regulatory regions is given in Fig. 2.

**Fig. 2.**
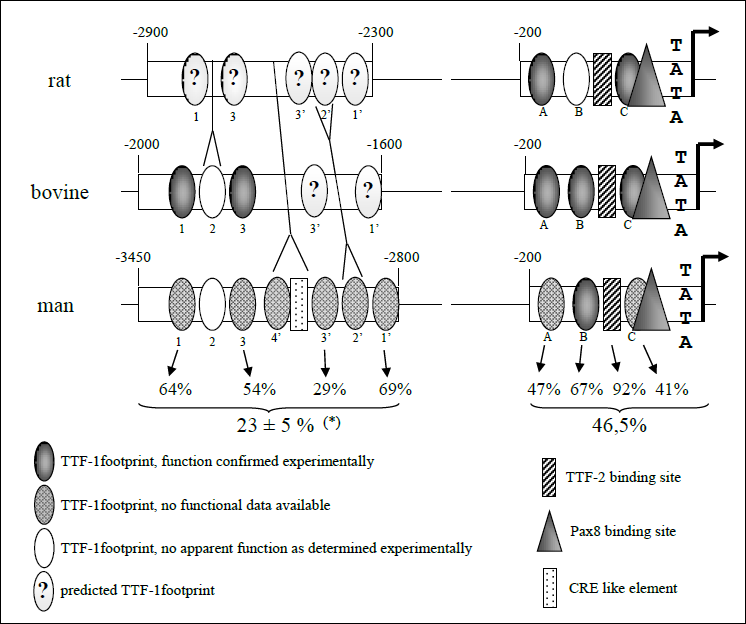
Schematic representation of available data on TTF-1 binding sites and their organization within rat, bovine and human Tg gene enhancer and promoter regions. Binding sites were named according to published data (TTF-1 binding sites 1 to 3: Christophe-Hobertus et al., 1992; TTF-1 binding sites 1’ to 4’: Berg et al., 1996; binding sites in the promoter: Civitareale et al., 1989). Functional data were extracted from Civitareale et al., 1989; Javaux et al., 1992; Donda et al., 1993; Christophe-Hobertus and Christophe, 1999a. Percentages of DNA sequence identity are given for binding sites conserved in all three species (note that TTF-1 and Pax8 binding overlap each other at site C). Overall percentages of DNA sequence conservation are also indicated for both promoter and enhancer regions (in this latter case, due to the large differences in size between species, a mean value ± half-range is given).

A 575bp-long fragment of rat genomic DNA, corresponding to the sequence of the DNaseI-hypersensitive enhancer region identified initially in the bovine, and thus encompassing the conserved TTF-1 binding sites doublet (see fig. 1), was amplified by PCR and cloned in front of reporter genes containing either the rat Tg gene proximal promoter or the SV40 promoter. As illustrated in fig. 3, transient transfection experiments performed in rat thyroid PCCl3 cells showed that the cloned sequences dramatically enhanced the activity of the rat Tg gene proximal promoter and significantly increased the activity of the heterologous SV40 promoter, indicating that the cloned fragment exhibited the activity of a transcriptional enhancer.

**Fig. 3.**
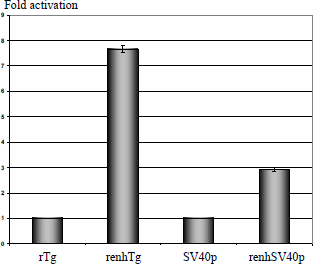
Functional characterization of the rat Tg gene upstream enhancer region. Luciferase reporter constructs were transiently expressed in thyroid PCCl3 cells. Measured Luc activities were normalized according to RLuc production from co-transfected pRLSV40 plasmid. Results are expressed as fold activation relative to prTgLuc (rTg) for prenhTgLuc (renhTg) or pGL3promoter (SV40p) for prenhTgSV40Luc (renhSV40p). The results shown are means ± SD from three and seven independent experiments respectively.

## Materials and methods

### Bioinformatics

DNA sequences alignment was performed and analyzed using DIALIGN 2.2.1 at BiBiServ (Morgenstern, 2004). EMBL/GenBank database accession numbers of the Tg promoter sequences are as follows; bovine: M35823; cat: AK071756; chimp: NW_001240369; dog: S61184; giraffe: AY007707; man: AF230667; mouse: NT_039621; rat: NW_047779.

### DNA constructions

Oligonucleotides used for PCR, DNA sequencing and EMSA experiments were purchased from Eurogentec (Seraing, Belgium).

All PCR-amplified sequences were verified by DNA sequencing in order to exclude the presence of sequence changes introduced during the PCR step.

The sequences of the 5’end of the rat Tg gene extending from position −180 to +9 (relative to transcription start as +1) were amplified by PCR from plasmid pTACAT3 (Sinclair et al., 1990; this plasmid was kindly given to us by Prof. R. Di Lauro, Stazione Zoologica A. Dohrn; Naples, Italy), flanked by XhoI and Hind III restriction sites in 5’ and 3’ respectively, and inserted between the corresponding cloning sites in pGL3basic (Promega Corp., Madison, Wisconsin, USA), yielding prTgLuc. The sequences of the 5’end of the rat Tg gene extending from position −2903 to −2329 (relative to transcription start as +1) were amplified by PCR from genomic DNA of rat thyroid PCCl3 cell line, flanked by Kpnl and Xhol restriction sites in 5’ and 3’ respectively; and inserted between the corresponding cloning sites in prTgLuc, yielding prenhTgLuc. The same PCR fragment was also inserted between the KpnI and XhoI cloning sites in pGL3promoter (containing the SV40 promoter; Promega Corp., Madison, Wisconsin, USA), yielding prenhTgSV40Luc.

### Cell culture and DNA transfections

PCCl3 cells were cultured as described by Fusco and co-workers (1987). The cells were trypsinized and seeded at 50% confluency the day preceding transfection. Transfections were performed using the FuGENE 6 transfection reagent as recommended by the supplier (Roche Diagnostics GmbH, Mannheim, Germany) at a usual ratio of FuGENE 6 (μL) to DNA (μg) of 3 to 1. To each individual cell culture dish (diameter: 3.5 cm) were added 500 ng of firefly luciferase reporter construct and 50 ng of pRLSV40 (Promega Corp., Madison, Wisconsin, USA), containing the renilla luciferase gene under the control of the SV40 enhancer-promoter and used as internal control for normalization in regard to transfection efficiency.

### Luciferase assays

Firefly and renilla luciferase activities were measured between 24 and 48 hours after transfection using the Dual-Luciferase^®^ Reporter Assay System as recommended by the manufacturer (Promega Corp.) and a TD-20/20 luminometer (Turner Designs, Sunnyvale, California, USA). All transfections were performed in duplicate within each individual experiment. For each reporter construct, the mean ratio (from the duplicates) of firefly luciferase activity to renilla luciferase activity was used as normalized luciferase activity.

